# Calcium transport by the *Mycobacterium tuberculosis* PE15/PPE20 proteins

**DOI:** 10.1101/2022.05.30.494056

**Authors:** Vishant Boradia, Andrew Frando, Christoph Grundner

## Abstract

Many aspects of eukaryotic physiology are regulated by calcium ions (Ca^2+^). Whether bacteria have similar Ca^2+^ systems for transport, storage, binding, and response to Ca^2+^ is not well understood. To identify components of Ca^2+^ signaling in *Mycobacterium tuberculosis*, we determined its transcriptional response to Ca^2+^. Overall, only few genes changed expression, suggesting a limited role of Ca^2+^ as a transcriptional regulator. However, two of the most strongly downregulated genes were the *pe15* and *ppe20* genes that code for members of a large family of proteins that localizes to the outer membrane. PE15 and PPE20 formed a complex and PPE20 directly bound Ca^2+^. Ca^2+^-associated phenotypes such as an increase in ATP consumption and increase in biofilm formation were reversed in a *pe15/ppe20* knockout strain, suggesting a direct role in Ca^2+^ homeostasis. To test whether the complex has a role in Ca^2+^ transport across the outer membrane, we created a fluorescence resonance energy transfer (FRET)-based Ca^2+^ reporter strain. A *pe15/ppe20* knockout in the FRET background showed a specific and selective loss of Ca^2+^ influx that was dependent on the presence of an intact outer cell wall. These data show that PE15/PPE20 form a Ca^2+^-binding protein complex that selectively imports Ca^2+^ and support the emerging idea of a general family-wide role of PE/PPE proteins in transport across the outer membrane.

## INTRODUCTION

Second messengers are a class of signaling molecules that permit a fast response and swift amplification of signals intracellularly. Among second messengers, Ca^2+^ is a particularly versatile signal in eukaryotes [1]. The functions of hundreds of human proteins are directly regulated by Ca^2+^, and almost every cellular process is affected by Ca^2+^ [1, 2]. Ca^2+^ signaling is initially facilitated by Ca^2+^ flux along membrane Ca^2+^ gradients that are carefully maintained by pumps and transporters.

Several bacteria also maintain a similar Ca^2+^ gradient between the inside and outside of the cell [3]. What’s more, Ca^2+^ has anecdotally been linked to bacterial processes such as motility, spore formation, gene expression [4, 5] and, in the case of *Yersinia*, also to virulence [6], suggesting a common bacterial Ca^2+^ sense-and-response system. However, while some components of Ca^2+^ signaling have been identified in bacteria, their number remains small. Ca^2+^ transporters have been annotated but few experimentally tested. Similarly, Ca^2+^ binding proteins have been predicted, but few experimentally confirmed. As a result, Ca^2+^ remains a poorly understood signaling mechanism in bacteria, and many fundamental questions about Ca^2+^ homeostasis remain unanswered, from the triggers of Ca^2+^ influx, the Ca^2+^-binding proteins, to the eventual cellular outcomes.

*Mycobacterium tuberculosis* (*Mtb*) is surrounded by a highly impermeable outer cell wall that is composed primarily of the complex phthiocerol dimycocerosates (PDIM) that form an outer membrane [7], creating a structure not unlike that of the outer membrane of gram-negative bacteria. While transport through the outer membrane in gramnegative bacteria is facilitated by characteristic beta barrel porins [8], no equivalent porins have been identified in *Mtb*. The impermeable outer membrane and lack of porins raise the question how *Mtb* transports small molecules such as nutrients, metabolites, but also Ca^2+^ across the outer membrane. In this way, the very first step in *Mtb* Ca^2+^ signaling remains unknown.

The mycobacterial PE/PPE proteins have long been a puzzle. They are predominantly found in pathogenic mycobacteria, where they take up a substantial share of the coding capacity [9], and many are substrates of a type VII secretion system [10, 11]. The PE/PPE proteins are associated with the outer membrane of the mycobacterial cell wall [12, 13]. Many PE/PPE proteins have been implicated in aspects of tuberculosis pathogenesis, but the molecular mechanisms have not been conclusively identified [14]. A recent milestone study showed that several PE/PPE protein pairs function as channels for nutrient transport across the outer mycobacterial membrane [15], defining an idiosyncratic transport system and suggesting a new and perhaps shared family-wide function for the PE/PPE proteins as small molecule transporters.

Here, we sought to further explore Ca^2+^-mediated processes in *Mycobacterium tuberculosis*. We identified regulation of ATP levels and biofilm formation by Ca^2+^. The transcriptional response to Ca^2+^ was narrow, and the most highly regulated genes were *pe15* and *ppe20*. We show that PE15/PPE20 form a complex, directly bind Ca^2+^, and facilitate the influx of Ca^2+^ into *Mtb* across the outer membrane. These data point to a functional Ca^2+^ sense-and-response system, identify physiologic processes regulated by Ca^2+^ and identify a new, specific PE/PPE Ca^2+^ import system across the outer membrane.

## RESULTS

### Ca^2+^ affects ATP levels and biofilm formation

The role of Ca^2+^ in *Mtb* physiology is almost entirely unknown. We initially tested for parallels with Ca^2+^ effects on other bacteria. Ca^2+^ transport in *E. coli* depends on ATP, and Ca^2+^ in turn increases intracellular ATP levels [16]. To test if Ca^2+^ has a similar effect on ATP levels in *Mtb*, we exposed *Mtb* to increasing concentrations of extracellular Ca^2+^ and measured the intracellular ATP levels. ATP levels increased by >2.5-fold in the presence of 1mM Ca^2+^ and as much as >4-fold in the presence of 10mM Ca^2+^ (Fig. 1A). These values are larger than those observed for *E. coli*, where 10mM Ca^2+^ resulted in 30% elevation in ATP [16]. To test the reverse effect, we cultured *Mtb* in presence of EGTA, a Ca^2+^-specific chelator and observed a reduction in ATP levels by ~50% (Fig. 1A).

**Figure 1:**
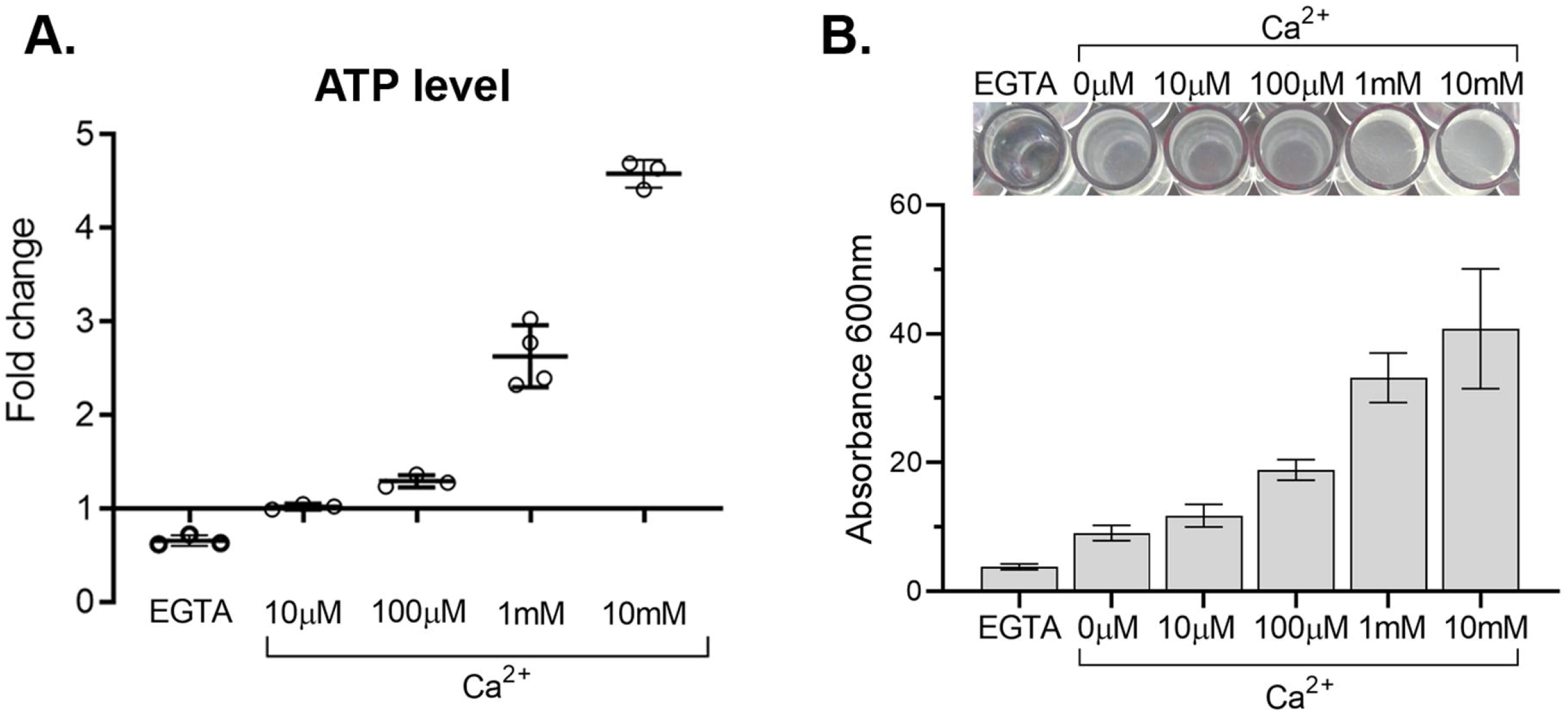
Ca^2+^ affects *Mtb* cellular processes. **A.** Effect of Ca^2+^ on cellular ATP levels. *Mtb* was grown in 48-well plates in detergent-free CTSM containing increasing concentration of Ca^2+^ (10μM – 10mM) and 1mM EGTA at 37 °C, treated with 0.1% Tween-80 and ATP was quantified using BacTitre-Glo reagent. Fold change was calculated by comparing to the Ca^2+^-free condition. Data represented are from biological triplicates. **B.** Ca^2+^ promotes biofilm formation. Biofilms were grown in the presence of 2+ increasing concentrations of Ca^2+^ (10μM – 10mM) and 1mM EGTA and were quantified using crystal violet. The experiment was repeated three times and a picture of a representative experiment is shown.

In some bacteria, a link between biofilm formation and Ca^2+^ has been proposed [17, 18]. To test if biofilm formation in *Mtb* is also dependent on Ca^2+^, we measured the generation of biofilm *in vitro* in response to increasing concentrations of Ca^2+^. Indeed, 1mM Ca^2+^ led to a 4-fold increase in biofilm when compared to low Ca^2+^ conditions (Fig. 1B). To rule out that this increase is simply a result of increased growth and thus biomass in high Ca^2+^ conditions, we tested for differences in growth of *Mtb* in the same Ca^2+^ concentrations as above. We detected no differences in growth (Fig. S1), showing that the biofilm effects are specifically due to biofilm generation, not to differences in biomass.

### *pe15/ppe20* transcripts are downregulated in response to Ca^2+^

In eukaryotes and in some bacteria, Ca^2+^ affects the transcription of a large number of genes [19]. To test for transcriptional effects of Ca^2+^ in *Mtb* and to identify genes potentially involved in Ca^2+^ homeostasis, we grew *Mtb in vitro* with and without 1mM Ca^2+^ and determined transcriptional effects by RNA-seq. At early timepoints, few transcripts changed abundance in response to Ca^2+^, although we detected a larger response after 3 days of Ca^2+^ exposure. Interestingly, two genes with reduced transcript abundance were apparent as early as day one and were the most downregulated genes by day three: *pe15* (Rv1386) and *ppe20* (Rv1387) (Fig. 2). This behavior in response to Ca^2+^ was reminiscent of metal transporter genes that are often regulated in response to changing metal concentrations, with importers typically down- and exporters typically upregulated in high metal concentrations.

**Figure 2:**
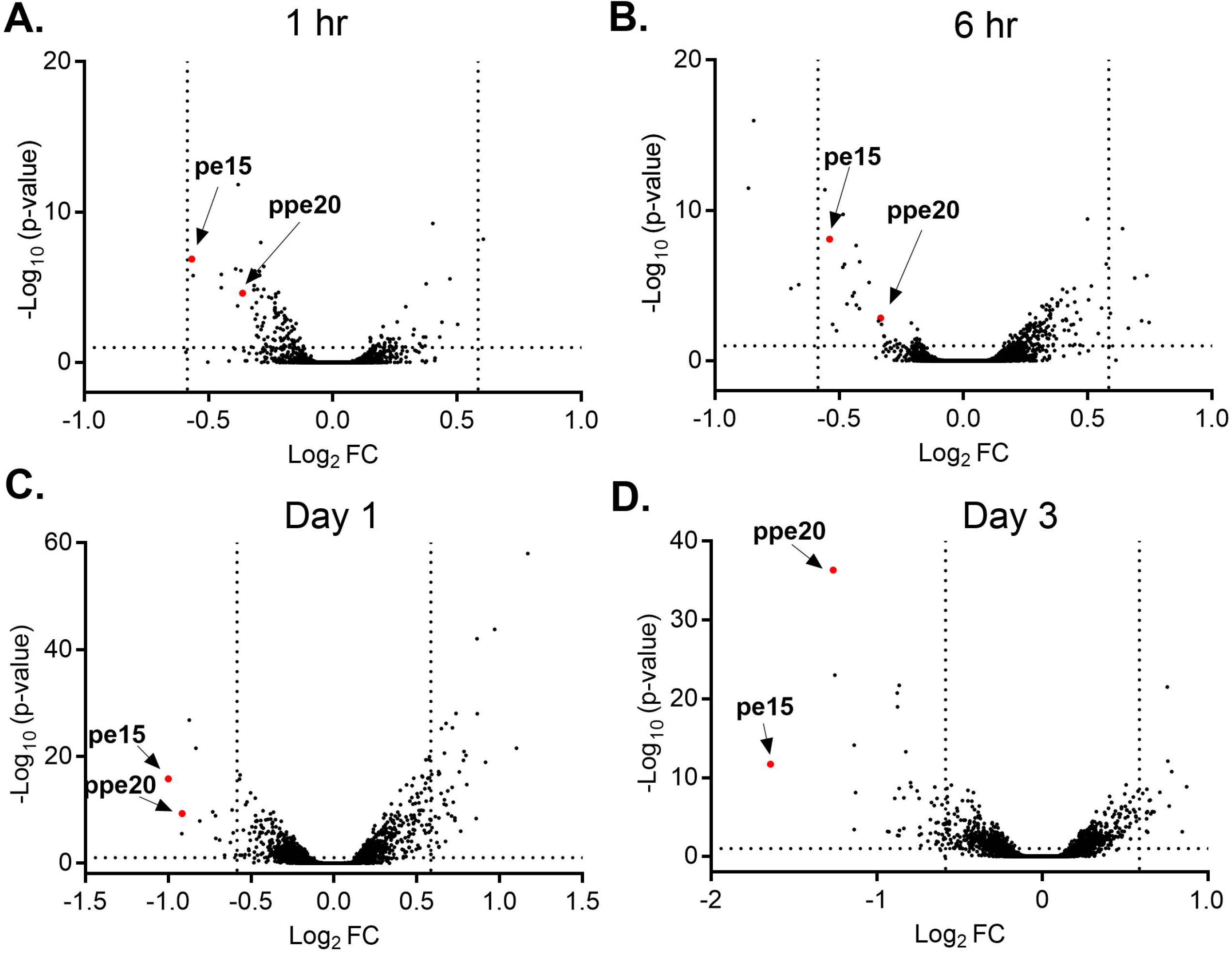
Ca^2+^ elicits a limited *Mtb* transcriptional response but downregulates *pe15* and *ppe20* transcripts. RNA-seq analysis of *Mtb* transcripts after exposure to Ca^2+^ for **A.** 1 hr **B.** 6 hr **C.** 1 day **D.** 3 days. Volcano plot shows downregulation of *pe15* and *ppe20* transcripts after exposure to 1mM Ca^2+^ after 1 and 3 days. Few other genes are significantly changed.

### PE15/PPE20 form a complex and bind Ca^2+^

Several PE/PPE proteins have been shown to form protein complexes, typically those coding in the same operon [20]. The *pe15* and *ppe20* genes are also operonic, suggesting that they could be a functional protein pair. According to previously published data, recombinant expression of PE15 and PPE20 individually in *E. coli* failed to produce any protein [20]. We could readily obtain recombinant protein when the two were expressed together, indicating a potential interaction. To conclusively show a PE15 and PPE20 interaction, we tested for an association by reciprocal pulldowns with tagged proteins. We co-expressed His-tagged PE15 and Strep II-tagged PPE20 from a dual expression plasmid and precipitated separately with beads against each tag. Both proteins were efficiently pulled down by either beads, indicating binding between PE15 and PPE20 (Fig. 3A). To further test the interaction, we co-expressed both proteins and visualized them by native PAGE. Both proteins migrated together, as shown by imaging for the respective tags, further confirming that they form a complex (Fig. 3B).

**Figure 3:**
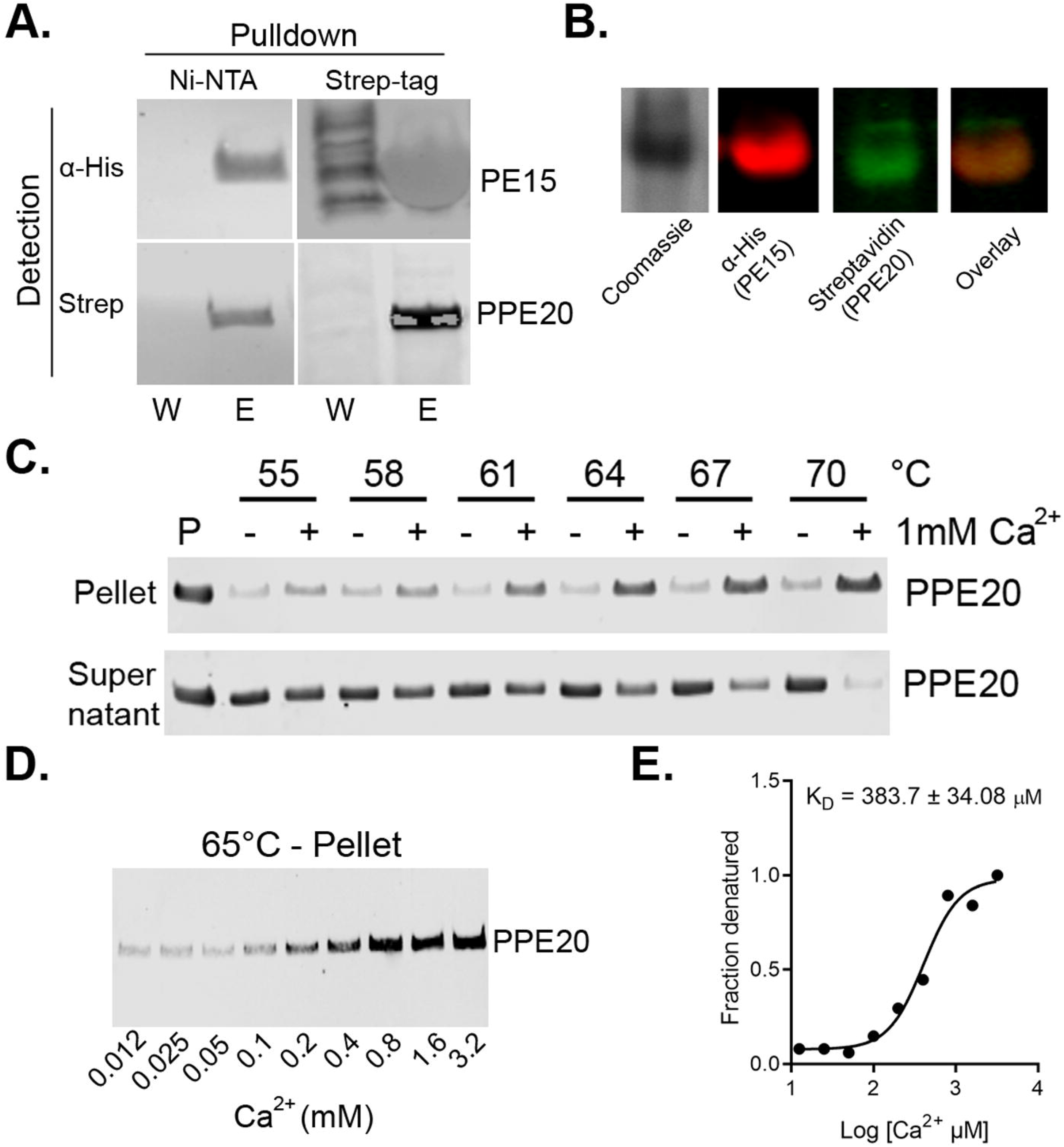
PE15 and PPE20 form a complex and PPE20 binds Ca^2+^. **A.** Western blot of PE15 with an N-terminal His tag and PPE20 with a C-terminal Strep tag shows the two co-purify. W = wash, E = elution **B**. PE15 and PPE20 migrate together on a native PAGE gel, also indicating complex formation **C.** Thermal melting experiment with Western blot readout shows differential thermal stability of PPE20 in the absence and presence of Ca^2+^. **D.** PPE20 stability is dose-dependent. **E.** Band intensities in D. plotted to estimate the K_D_ of PPE20 for Ca^2+^ (383μM).

To test whether PE15 and/or PPE20 directly bind Ca^2+^, we incubated recombinantly expressed protein with Ca^2+^ and tested for protein stability using thermal denaturation and a gel readout (Fig. 3C). After heating and precipitation, PPE20 showed clear differential thermal stability when incubated with Ca^2+^, indicating Ca^2+^ binding. By using different concentrations of Ca^2+^ in the same assay, we determined a Ca^2+^ denaturation curve and estimated a K_*D*_ of 383.7 ± 34.08 μM (Fig. 3D,E). Interestingly, in a departure from typical ligand-induced thermal changes, Ca^2+^ binding decreased thermal stability of PPE20. PE15 did not produce interpretable results in this assay. Ligand binding in the context of transport suggests that the PE15/PPE20 complex may be a specific channel, as channels typically bind to their ligands [21].

### PE15/PPE20 KO reverses Ca^2+^-dependent phenotypes

To test for a phenotypic link of the PE15/PPE20 protein complex in Ca^2+^-dependent processes, we generated an *Mtb pe15/ppe20* knockout (KO) strain by recombineering [22]. We next tested the *pe15/ppe20* KO strain for altered ATP levels. The KO showed reduced Ca^2+^-dependent increase in ATP production which was reverted to wild type (WT) levels by complementation with *pe15/ppe20* (Fig.4A). We next tested whether the PE15/PPE20 complex also affects Ca^2+^-dependent biofilm formation. The KO blocked the effect of Ca^2+^ on biofilm formation, and complementation with *pe15/ppe20* restored the phenotype (Fig. 4B). These data show that the PE15/PPE20 complex is not only transcriptionally responsive to, but regulates functions related to Ca^2+^.

**Figure 4:**
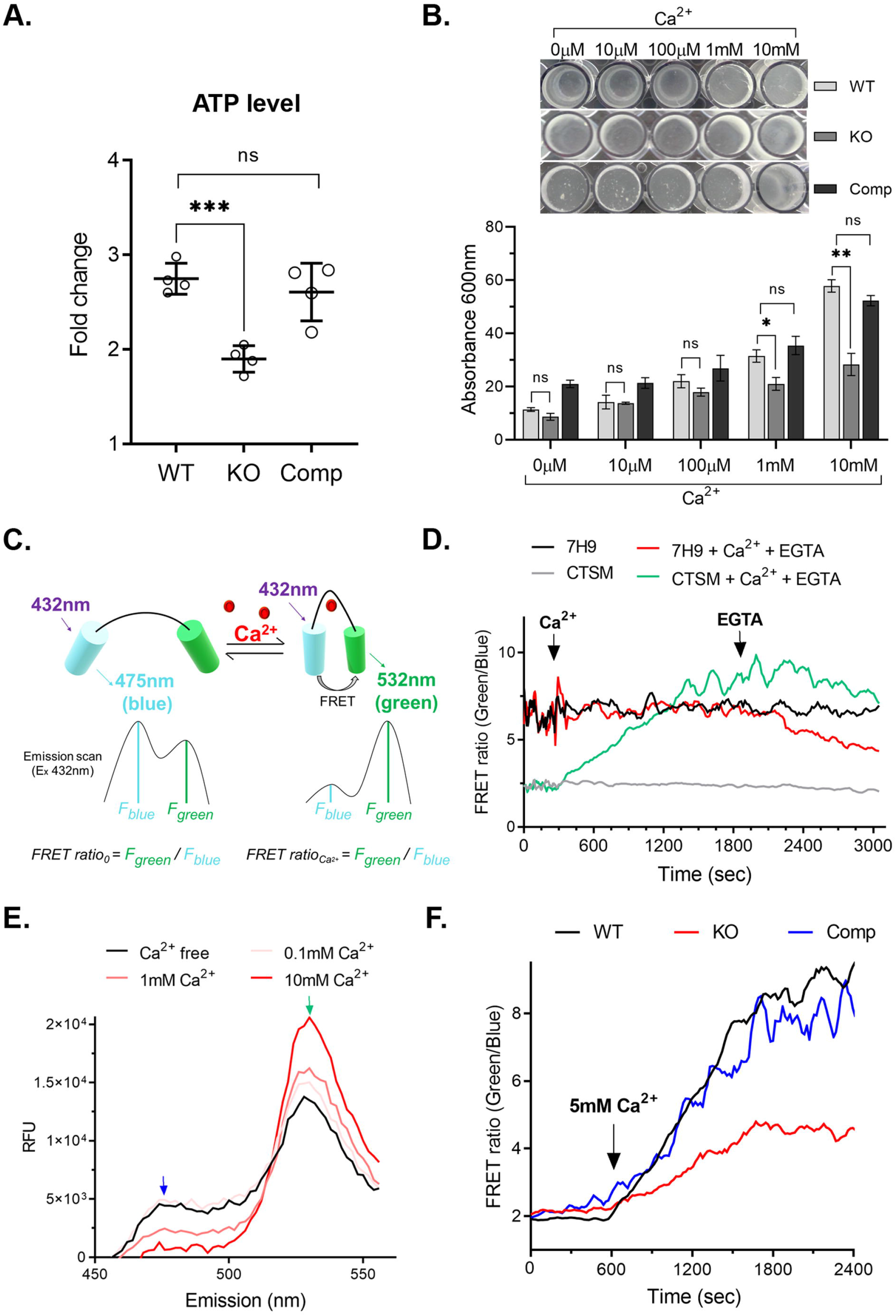
*pe15/ppe20* affects Ca^2+^ associated processes and facilitates Ca^2+^ uptake. **A.** *Mtb* was treated with 1mM Ca^2+^ at 37°C and ATP was quantified using bactitreGlo reagent. Fold change was calculated by comparing to the Ca^2+^-free condition. Data shown are from four experiments (^ns^P >0.05, ***P <0.001). **B.** Cultures were grown in the presence of increasing concentrations of Ca^2+^ or EGTA and biofilm was quantified by crystal violet. The experiment was repeated three times (^ns^P >0.05, *P <0.05, **P <0.01). **C.** Schematic of the Ca^2+^ FRET probe and its ratiometric signal. **D.** Ca^2+^ FRET signals in different media show that even low concentrations of Ca^2+^ in standard 7H9 compromise Ca^2+^ detection. FRET ratio was calculated by calculating the green:blue fluorescence ratio and was plotted against time. **E.** The FRET probe generates a robust, dose-dependent FRET signal. Emission scan of *Mtb-twitch* incubated with increasing concentrations of Ca^2+^ for 30mins at 37°C. **F.** FRET trace over time shows reduced Ca^2+^ influx in the *pe15/ppe20* KO strain.

### The PE15/PPE20 complex is a Ca^2+^ importer

A recent study showed selective porin function of PE/PPE proteins [15], and the decrease of transcript levels of *pe15/ppe20* in the presence of Ca^2+^ was consistent with the behavior of a Ca^2+^ channel. To directly test the idea that PE15/PPE20 has a role in Ca^2+^ transport, we created an *Mtb* reporter strain using a fluorescence resonance energy transfer (FRET) system that detects intracellular Ca^2+^, Twitch [23]. Twitch is based on a minimal Ca^2+^ binding domain from troponin C that is optimized for maximal Ca^2+^ selectivity and ratiometric signal [23] (Fig. 4C). We ectopically expressed Twitch-2B in the H37Rv background (*Mtb*-twitch) and tested for Twitch expression in different cell fractions. Twitch was expressed only in the cytoplasm, indicating that it would report only on cytoplasmic Ca^2+^ (Fig. S2A). We could only obtain a robust FRET signal in Ca^2+^-free Sauton’s medium, not 7H9 medium which contains low amounts of Ca^2+^ (Fig. 4D). We thus used Sauton’s medium for all Ca^2+^ measurements. To further test the Twitch reporter in *Mtb*, we measured the Ca^2+^ signal at different Ca^2+^ concentrations. The FRET signal was robust, dose-dependent, and comparable to that of Twitch in previously described non-bacterial systems [23] (Fig. 4E). We next tested the selectivity of the probe over the closest earth alkali metal neighbor, Mg^2+^. Mg^2+^ did not generate a FRET signal (Fig. S2B). These data show that Twitch is a sensitive probe for measuring *Mtb* intracellular Ca^2+^ continuously, establish conditions for detecting intracellular Ca^2+^ changes, and show that *Mtb* readily takes up extracellular Ca^2+^. These data reveal a similar *Mtb* response to extracellular Ca^2+^ to that previously observed in *E. coli* [24].

To directly test for a role of the PE15/PPE20 complex in Ca^2+^ transport, we measured Ca^2+^ uptake in the *Mtb*-twitch strain and a strain expressing *Mtb*-twitch in the *pe15*/*ppe20* KO background. The WT produced a robust FRET signal as before upon addition of 5mM Ca^2+^. The KO, however, showed reduced influx, consistent with a selective import channel (Fig. 4F). Complementation of the KO with *pe15/ppe20* expressed from an extrachromosomal plasmid restored Ca^2+^ levels to those in WT. To test if PE15/PPE20 localize to the outer membrane as has been shown for other PE/PPE pairs [15, 25], we tested for PPE20 expression in different cell fractions. PPE20 was only detected in the cell wall fraction (Fig. S3A). To further test if PE15/PPE20’s transport role is associated with the cell wall/outer membrane, we permeabilized the *Mtb* cell wall by treatment with lysozyme and Triton-X100 [26]. The difference in Ca^2+^ influx between WT and KO was reduced by permeabilization, indicating that PE15/PPE20 facilitate transport across the outer membrane (Fig.S3B). To test for selectivity of the PE15/PPE20 complex for Ca^2+^, we grew the WT and KO strains in low Mg^2+^ conditions. Because Mg^2+^ is essential for *Mtb* growth [27], an effect of PE15/PPE20 on Mg^2+^ transport would be detectable as a growth defect. WT and KO grew equally (Fig S3C, D), indicating that the PE15/PPE20 complex is specific for Ca^2+^ even over other earth alkali metals.

## DISCUSSION

Ca^2+^ signaling is ubiquitous in eukaryotes [1] but in bacteria, only few components and functions of Ca^2+^ signaling have been characterized. In *Mtb*, the cell wall generally prevents passage of charged solutes, presenting a hurdle for Ca^2+^ uptake not encountered in most bacteria. In the absence of typical porins, the transport processes that allow for transfer through the cell wall and outer membrane have long been unknown. Here, we identify a specific Ca^2+^ channel that consists of PE15 and PPE20. In addition, we identify several phenotypes associated with elevated Ca^2+^ in *Mtb*: Ca^2+^ leads to an increase in cellular ATP concentrations, an effect that has also been reported in *E. coli* and was suggested to be a mechanism to sustain the increased activity of Ca^2+^ ATPase efflux pumps required to reset Ca^2+^ levels [16]. We also show a clear contribution of Ca^2+^ to biofilm formation. The role of biofilms for *Mtb* pathogenesis and tuberculosis treatment has long been unclear, but recent evidence supports the presence of biofilms in infected lungs of nonhuman primates and human patients and a role in *Mtb* pathogenesis and drug susceptibility [28].

The PE/PPE proteins have long been the subject of much speculation. They are specific not only to mycobacteria but are predominantly found in pathogenic or slow-growing mycobacteria. Although they make up ~10% of *Mtb’s* genetic coding potential, their function has long remained unclear [14]. Their large number and sequence variation is reminiscent of variable surface proteins that serve as antigenic decoys in other pathogens. However, phylogenetic analyses showed that sequence variation does not arise through immunogenic pressure and does not support this idea [14]. In contrast, a recent hallmark study showed channel-like function of PE/PPE complexes specifically transporting several carbon sources, Mg^2+^, and phosphate across the outer mycobacterial cell membrane [15]. We now show a PE/PPE complex with such a channel-like function for Ca^2+^. What’s more, we show direct binding of PPE20 to the cargo, a behavior that is more characteristic of specific channels than porins. The selectivity of PE15/PPE20 for Ca^2+^ over even the closely related Mg^2+^ further argues for a specific channel rather than a porin, which often show more indiscriminate transport [29]. In fact, another PE/PPE pair, PE20/PPE31 specifically transports Mg^2+^ [15]. Interestingly, several other PE-PGRS proteins bind Ca^2+^ [30], could also contribute to Ca^2+^ import, and could explain the residual Ca^2+^ import seen in the *pe15/ppe20* KO strain. A link of PE15/PPE20 to pathogenicity is highly plausible given the association of PE/PPE proteins with the outer mycobacterial cell membrane [31], the presence of PE/PPE proteins primarily in pathogenic mycobacteria, their association with the type VII secretion system, and previous studies directly implicating other PE/PPE proteins in virulence [14]. The source of Ca^2+^ for the PE/PPE channel is arguably host Ca^2+^. In this way, *Mtb* may be able to eavesdrop on the many host cell signaling events that involve Ca^2+^, for example phagocytosis, which is Ca^2+^-dependent [32] and marks the beginning of *Mtb’s* intracellular stage.

How do PE/PPE proteins facilitate transport? Crystal structures show that these proteins, unlike classical porins, fold into long alpha helices, with no structural information available for the long repetitive C-terminal sequences of the PPE proteins that are thought to be intrinsically disordered. A potential model for how the alpha helical sections of PE/PPE proteins may form pores was recently suggested by structures of the ESX type VII secretion system component EspB [33], which is a naturally occurring fusion of a PE and a PPE protein that forms a donut-shaped heptamer with a large central pore. Although the helices that potentially traverse the cell membrane did not have the requisite outward-facing hydrophobic residues, other models in which the PPE C-termini provide a hydrophobic sheath or association with other membrane-spanning proteins of the ESX complex are possibilities. In our studies, Ca^2+^ binding led to an atypical increase in denaturation of PPE20. This change likely indicates a local unfolding event. The precise nature of this unfolding event and how it impacts transport remains to be identified along with the larger transport mechanics of the PE/PPE proteins. Although our data show that PE15/PPE20 are necessary for efficient Ca^2+^ uptake, they may not be sufficient. The genetic and functional association of the PE/PPE proteins with the ESX secretion system suggests that other ESX components such as an ATPase could be recruited for PE/PPE transport functions.

The PE and PPE proteins can be further stratified into subfamilies by C-terminal sequence motifs [14] that perhaps also indicate distinct functions. While all currently known PE/PPE channels including PE15 contain a minimal PE-only protein, the PPE proteins identified as components of transporters prior to our study belong to the PPE-SVP subfamily. However, PPE20 belongs to the PPE-PPW subgroup and is the first example of a PE/PPE channel outside of the PPE-SVP group. Similarly, the previous PE/PPE transporter pairs are associated with the ESX5 secretion system [34], while PE15/PPE20 is the first system associated with ESX3 [31]. Together, our data support the emerging theme that the PE/PPE protein family is a mycobacterial transporter family that solves the conundrum of an outer cell membrane so impermeable as to exclude vital nutrients. The PE/PPE channels appear to be highly specific, which could explain their large number, as *Mtb* requires access to many different nutrients without compromising the outer membrane’s barrier function. The full set of PE/PPE transporter components remains to be identified, as does the generality of a transport function across all PE/PPE subfamilies and their specific cargoes. However, these studies could explain the widely different phenotypes associated with these idiosyncratic proteins and eventually explain their diverse roles in physiology and pathogenesis.

## METHODS

### Media and growth conditions

*M. tuberculosis H37Rv* was used as a parental strain for generation of all mutants and were cultured at 37 °C in either Middlebrook 7H9 medium (Difco) with 10% (vol/vol) oleic acid-albumin-dextrose-catalase (OADC) enrichment (BBL; Becton Dickinson), 0.5% glycerol, and 0.05% Tween 80 (referred to as “7H9+GOT”) or 7H10 agar supplemented with 10% OADC and 0.5% glycerol or Chelex treated Sauton’s medium (CTSM) with additional supplements as indicated. Chelex treated Sauton’s medium (referred to as “CTSM”) consisted of 0.5 g KH_2_PO_4_, 4g L-asparagine, 2 g citric acid, 6% glycerol, adjusted to pH ~7.0, and treated overnight with chelex-100 resin (10 g/L) (sigma) to remove trace metal ion contaminants including calcium. After filtration, 0.5 g/L of MgSO_4_, 0.05 g/L ferric ammonium citrate, 0.1ml of 1% zinc sulfate and 0.05% tween 80 was added. The pH was adjusted to 6.9 and the medium was sterilized by filtration. Strains bearing antibiotic cassettes were cultured with 50 μg/mL hygromycin or 30 μg/mL kanamycin or 25 μg/ml zeocin as appropriate.

### Biofilm formation

*M. tuberculosis* biofilms were generated and quantified according to a published protocol [35] in 48-well polystyrene plates. Briefly, cells were grown to an OD_600_ of 0.8-1 in 7H9+GOT medium, washed twice with CTSM and diluted to an OD_600_ of 0.01 in CTSM without detergent and added to each well supplemented with varying concentrations of Ca^2+^. Outer wells were filled with water and the plate was incubated at 37°C for 4 weeks without shaking. The plates were photographed and the processed for crystal violet staining. The medium was removed, biofilms were dried and incubated with 500μl of 1% crystal violet for 10min. Wells were washed three times with water and dried again. Absolute ethanol (1ml) was added to each well and incubated for 10min. Then 3-fold serial dilutions were read at A600 on a spectrophotometer in a 96-well plate. The represented bar graph is the average Absorbance 600nm readings of four biological replicates and Welch’s t test was applied to determine the significance (p-value).

### ATP measurement

*Mtb* cells from a 7H9+GOT medium culture (OD_600_~1) were washed twice and diluted to an OD_600_ of 0.01 in CTSM without detergent supplemented with different concentrations of CaCl_2_ or 1mM EGTA. Cultures were grown in a 48 well plate at 37°C for 14 days. Cells were treated with 0.1% Tween-80 overnight to create a homogenous suspension. ATP was quantified by incubating 50μl of the bacteria in triplicate with 50μl of the BacTiter-Glo™ reagent for 15mins followed by a luminescence reading. ATP production in the Ca^2+^ treated cultures were calculated as fold change compared to the Calcium free condition. The represented bar graph is the average fold change of four biological replicates and Welch’s t test was applied to determine the significance (p-value).

### RNA sequencing

*Mtb* H37Rv strain was grown to an OD_600_ of ~1 in 7H9+GOT medium. Cells were washed twice with CTSM and subculture in CTSM starting at an OD_600_ of 0.05 and was grown to an OD_600_ of ~1. The culture was again subcultured in CTSM starting at an OD_600_ of 0.05 and incubated at 37°C to an OD_600_ of 0.2. The cultures were supplemented with or without 1mM CaCl_2_ in triplicate. At the indicated times, cells were pelleted at 4000g for 5min at 4°C, resuspended in Trizol and lysed by bead beating for 30sec at 6m/s for three cycles with intermittent cooling on ice. Cell debris were pelleted at 20,000g for 1min, and the supernatant was transferred to a heavy phase lock gel tube containing 300 μl chloroform. The tubes were inverted for 2min and centrifuged at 20,000g for 5min. RNA in the aqueous phase was precipitated using 300 μl isopropanol and 300 μl high salt solution (0.8M Na citrate, 1.2M NaCl). RNA was purified using QIAGEN RNeasy kit and ribosomal RNA was depleted using the Ribo-Zero rRNA removal magnetic kit (Illumina). The cDNA library was generated using the NEBNext Ultra II RNA Library Prep Kit and each replicate was barcoded in the DNA library using the NEBNext Multiplex Oligos for Illumina. Libraries were quantified using the KAPA qPCR quantification kit, pooled, and sequenced at the University of Washington Northwest Genomics Center with the Illumina NextSeq 500 High Output v2 Kit. Read alignment was performed using the Bowtie 2 custom processing pipeline (https://github.com/robertdouglasmorrison/DuffyNGS, https://github.com/robertdouglasmorrison/DuffyTools). Gene expression changes were identified using a combination of five differential expression (DE) tools within DuffyTools. The 5DE tools included round robin, RankProduct, significance of microarrays (SAM), EdgeR, DeSeq2. Each DE tool measurement was combined using the weighted average of fold change and significance (p-value). Genes with averaged absolute fold change more than 1.5 fold and p-value <0.01 were considered differentially expressed.

### Cloning, co-expression, and purification of recombinant PE15 and PPE20 proteins from *E.coli*

pETDuet dual expression plasmid was used to co-express PE15 and PPE20. PE15 was cloned in the MCS-1 region with N-terminal His tag and PPE20 was cloned in the MCS-2 region with a C-terminal Strep II tag. *Mtb* Rv1386 (PE15) and Rv1387 (PPE20) genes were amplified from *Mtb* H37Rv genomic DNA using the primers Duet 1-5 which included Gibson overlap sequence (Table 1). The linearized pETDuet plasmid and the region between MCS-1 and MCS-2 were prepared by PCR amplification using the primers Duet 6,7 and Duet 8,9 respectively (Table 1). Finally, all the four purified PCR products i.e. PE15, PPE20, region between MCS-1/MCS-2 and linearized pETDuet plasmid were ligated using the Gibson Assembly® (NEB) to generate the pETDuet pe15/ppe20 plasmid. Cloning was confirmed by sequencing. The vector was transformed into *E. coli BL21*(*DE3*) and a single colony was picked and was grown at 37 °C in Terrific broth medium containing ampicillin. Protein expression was induced by addition of 0.5mM isopropyl-δ-d-thiogalactoside (IPTG) at OD_600_ of ~0.4, culture was then maintained at 16 °C for 16hrs. Cells were harvested, and pellets were processed for purification. For His-PE15 purification, pellets were resuspended in buffer A (50 mM Tris pH 8.0, 150 NaCl and 10% glycerol) containing 20mM imidazole and 1mM AEBSF and lysed by sonication. Lysate was centrifuged at 35,000g for 30min at 4°C and the supernatant was loaded on Ni-NTA column. Column was thoroughly washed with buffer A containing 20mM imidazole followed by elution with buffer A containing 250mM imidazole. For PPE20-strep II tag, pellets were resuspended in buffer A containing 1mM AEBSF and lysed by sonication. Lysate was centrifuged at 35,000g for 30min at 4°C and the supernatant was loaded on Strep-tactin column. Column was thoroughly washed with buffer A followed by elution with buffer A containing 2.5mM desthiobiotin. Both the purified proteins were immediately dialyzed against buffer A and stored at −80 °C. Interaction of pe15/ppe20 was confirmed by Western blot. The wash and elution fraction from the Ni-NTA and strep tag purification were loaded on either SDS-PAGE or Native-PAGE and the proteins were transferred onto a nitrocellulose membrane. Blots were probed with mouse α-His antibody followed by IRDye® 680RD Goat α-Mouse IgG Secondary Antibody (LI-COR) to detect the His tagged PE15, and were also probed with IRDye® 800CW Streptavidin to detect the PPE20-strep tag. The blots were scanned on LI-COR Odyssey platform using the 700nm and 800nm channel.

**Table 1:**
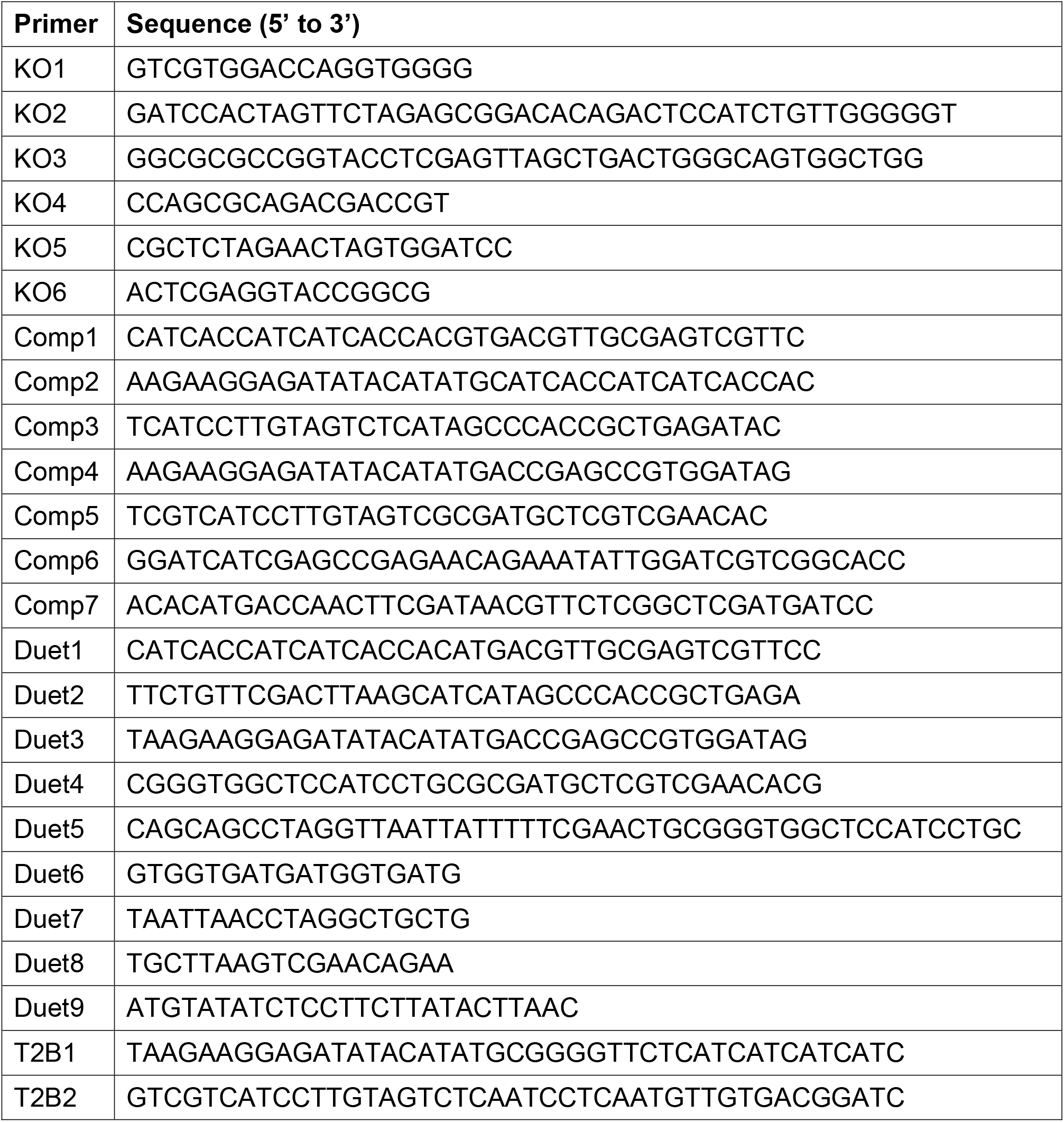
Primers used in this study.

### Thermal melting to determine Ca^2+^ binding to PPE20

Purified PPE20 protein was incubated with or without 1mM CaCl_2_ in a thermocycler for 5 min at 25°C followed by incubation at varying temperature (55-70°C) for 3mins. The reaction was immediately cooled to 4°C and was centrifuged at 20,000g for 30min at 4°C to separate the native (supernatant) and the denatured (pellet) fractions. Pellet fraction was solubilized by boiling with 2x SDS loading buffer. PPE20 in each of the supernatant and pellet fraction was detected by western blot using rabbit α-strep II tag antibody followed by IRDye® 800CW Goat α-Rabbit IgG Secondary Antibody. To determine the K_D_ value, PPE20 was incubated with varying concentration of CaCl_2_ (12.5μM – 3.2mM) for 5 min at 25°C, 3min at 65°C followed by incubation at 4 °C. PPE20 in the pellet fraction was detected by western blot as described above. The band intensities were calculated using the Image studio software and converted to fraction denatured relative to the total protein. The graph of fraction denatured v/s log Ca^2+^ concentration was plotted using GraphPad Prism 9.3 and the K_D_ value was determined by fitting a dose response curve using the least square regression method.

### Creation of *pe15/ppe20* knock out and complemented strain in *Mtb* H37Rv

The knockout strain was created by recombineering as described previously [22]. Initially 500bp upstream of pe15 (primers KO1 and KO2), 500bp downstream of ppe20 (primers KO3 and KO4) and hygromycin cassette (primers KO5 and KO6) were amplified separately (Table 1). Both PCR fragments were Gibson ligated to the 5’ and 3’ of a hygromycin-resistance cassette, respectively, to generate the recombineering cassette. This linear recombineering cassette was PCR amplified, purified and electroporated into *Mtb* H37Rv strain carrying recombineering plasmid pNIT [36]. Hygromycin positive colonies were screened, and the positive clone was confirmed by DNA sequencing. For the complemented strain, both *pe15* (His-tag) and *ppe20* (flag tag) genes were separately PCR-amplified using primers Comp 1-3 and Comp 4, 5, respectively (Table 1), and Gibson cloned in pDTCF plasmid (Zeocin) separately under an anhydrotetracycline (ATc)-inducible promoter. The ATc promoter along with PPE20 gene was amplified using primers Comp 6, 7 (Table 1) from pDTCF-PPE20 vector and Gibson cloned in pDTCF-PE15 vector downstream of pe15 gene. The resulting plasmid pDTCF-pe15/ppe20 carries both genes with individual ATc promoters. The plasmid was electroporated into the knockout strain and positive clones were selected by growth in hygromycin and zeocin. This strain expressed His-PE15 and PPE20-Flag proteins when induced with 100ng/ml ATc.

### Creation of *Mtb* Ca^2+^ FRET reporter strain

The Ca^2+^ FRET reporter Twitch-2B was PCR amplified from the plasmid Twitch-2B pRSETB obtained from Addgene [23] using the primers T2B1 and T2B2 (Table 1) which included Gibson overlap sequence. The PCR product was Gibson cloned in the *E. coli-mycobacterial* episomal shuttle vector pDTCF (kanamycin) under the anhydrotetracycline (ATc)-inducible promoter. The plasmid was electroporated into wild type *Mtb* H37Rv (WT), *pe15/ppe20* knockout (KO) and *pe15/ppe20* complemented strain (Comp), and the expression of Twitch was induced using 100nM ATc. The expression of Twitch was confirmed by measuring the fluorescence of uninduced and induced cells using a plate reader at three different wavelengths (432/475nm, 432/532nm and 488/532nm). The localization of Twitch in WT strain was confirmed by Western blot. Briefly, 100ml culture of uninduced and induced *Mtb-Twitch* was grown in 7H9/OADC/tween medium. Cells were washed twice in PBS and resuspended in lysis buffer (50mM Tris pH 7.4, 150mM NaCl) containing protease inhibitor cocktail. Cells were lysed by bead beating and different fractions i.e. cytosolic (CYT), cell membrane (CM), and cell wall (CW) were prepared according to published protocol [37]. Western blot was performed and the expression of Twitch protein (His-tagged) was detected by Mouse α-His antibody and the purity of cellular fraction was confirmed by Western blot using α-LAM (CM and CW fraction) and α-FtsZ (CYT) antibodies (obtained from BEI resources, NIAID, NIH).

### Ca^2+^ uptake assay

#### Media

*Mtb-twitch* was grown in either 7H9+GOT or CTSM to an OD_600_ of 1.0. Cells (50μl) were aliquoted in duplicate in a 384 well plate and fluorescence was immediately measured at wavelengths 432/475nm (Blue) and 432/532nm (Green) continuously for 5mins using the plate reader pre-set at 37°C. Ca^2+^ (5mM) followed by EGTA (5mM) was added and the fluorescence was again measured for 30min and 15mins. Fluorescence readings were corrected by subtracting the readings from wells containing uninduced culture. The FRET ratio was calculated as the green:blue fluorescence ratio and plotted against time.

#### Different Ca^2+^ concentrations

*Mtb-twitch* culture was grown in CTSM to an OD_600_ of 1.0. Cells (50μl) supplemented with different concentrations of Ca^2+^ were aliquoted in duplicate in a 384 well plate and incubated for 30min at 37°C. Emission scan (from Em450-550nm) using Ex432nm was recorded, and the relative fluorescence units (RFU) were corrected by subtracting the readings from uninduced culture wells.

#### Mutant strains

Wild type *Mtb* H37Rv (WT), pe15/ppe20 knockout (KO) and pe15/ppe20 complemented strains (Comp) expressing Twitch were grown to OD~1 in 7H9+GOT. Cells were washed twice with CTSM and subculture in CTSM with or without 100mM ATc starting at 0.05OD. Cells were grown to a density of OD_600_ of ~0.5, washed twice with CTSM and resuspended in CTSM at a dilution of 1 OD/ml. Cells (50μl) were aliquoted in duplicate in a 384 well plate and fluorescence was immediately measured at wavelengths 432/475nm (Blue) and 432/532nm (Green) continuously for 10min using the plate reader pre-set at 37°C. Calcium (5mM) was added to each well using multichannel pipette and the fluorescence was again measured for 30mins at 37°C. Fluorescence readings were corrected by subtracting the readings from uninduced culture wells. FRET ratio was calculated by calculating the green:blue fluorescence ratio and was plotted against time. Increase in FRET ratio would indicate increase in intracellular calcium concentration. The assay was repeated by treating cells with permeabilizing reagent [26] (0.1% Triton-X100 and 2μg/μl lysozyme) at time 0min.

## SUPPLEMENTARY FIGURE LEGENDS

**Supplementary Figure 1: Growth curves of *Mtb H37Rv*** in **A.** Chelex-treated Sauton’s medium (CTSM) and **B.** 7H9 medium containing glycerol, OADC and Tween-80 supplemented with different concentrations of CaCl_2_.

**Supplementary Figure 2: Intracellular Ca^2+^ detection by FRET in *Mtb-twitch*. A.** Western blot showing localization of Twitch-2B (His tagged) in the cytosolic fraction. LAM is a cell envelope marker and FtsZ is a cytosolic marker. **B.** The FRET signal is highly specific for Ca^2+^ over the congener Mg^2+^. The His-tagged Twitch protein was purified from the cytosolic fraction using Ni-NTA column, treated with EDTA and desalted. Emission scan of the purified protein was recorded upon incubation with either Ca^2+^ or Mg^2+^.

**Supplementary Figure 3: Transport by PE15/PPE20 is associated with the cell wall and is specific for Ca^2+^**

**A.** Western blot showing localization of PPE20 to the cell wall. Subcellular fractions from the complemented *pe15/ppe20* KO strain expressing PPE20-FLAG were probed with α-FLAG antibody. Same cell fractions as in Fig. S2B were used, see controls therein. **B.** WT, *pe15/ppe20* KO, and complemented strains were transformed with Twitch, permeabilized with lysozyme and Triton-X100, and Ca^2+^-FRET signal measured. (P) permeabilized. **C.** *Mtb* strains were grown in 96-well plates in detergent-free medium containing different concentrations of Mg^2+^ at 37°C and photographed after 14-days incubation to monitor growth. **D.** *Mtb* strains were grown in detergent-free medium containing 10μM Mg^2+^ at 37°C for 14 days, treated with 0.1% Tween-80 and growth was quantified by alamarBlue assay. Data shown are from technical triplicates.

