## Supplementary figures and images for "Calcium transport by the *Mycobacterium tuberculosis* PE15/PPE20 proteins"

### Supplementary Figure S1

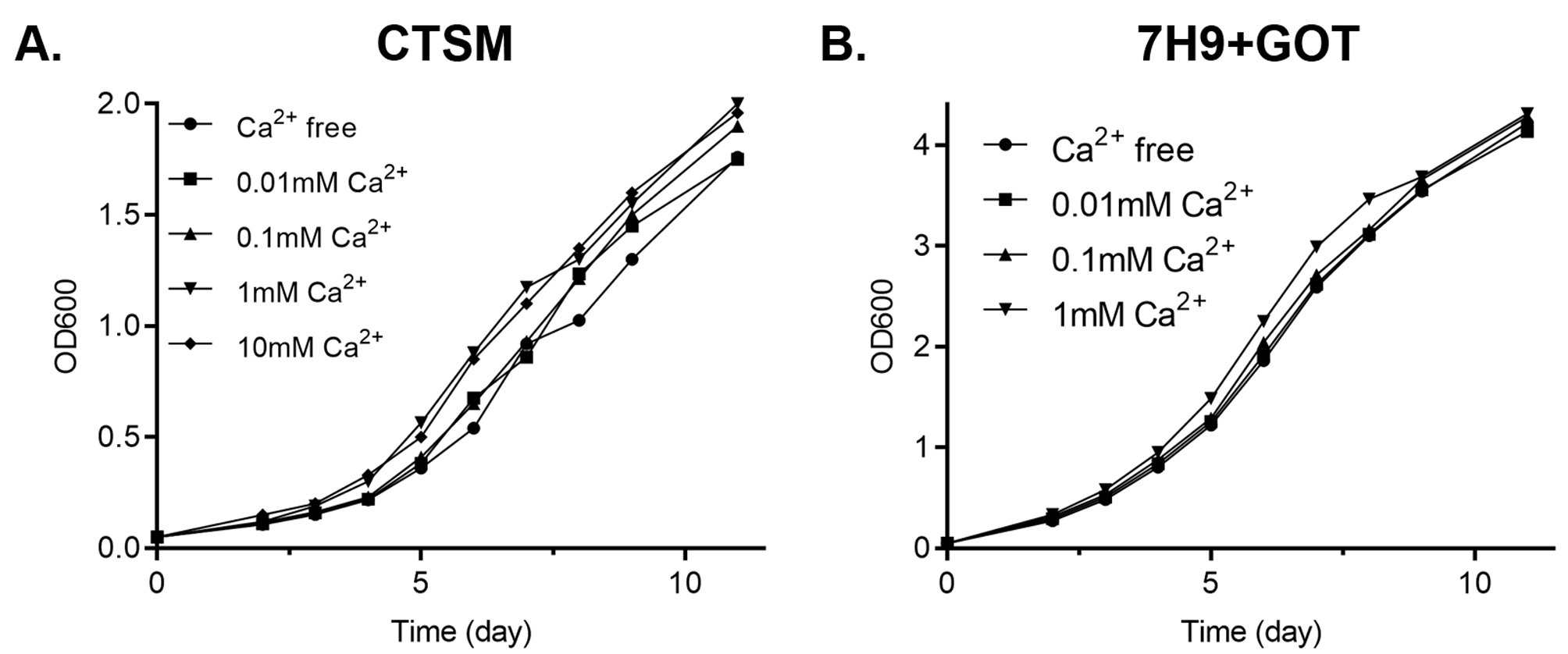

### Supplementary Figure S2

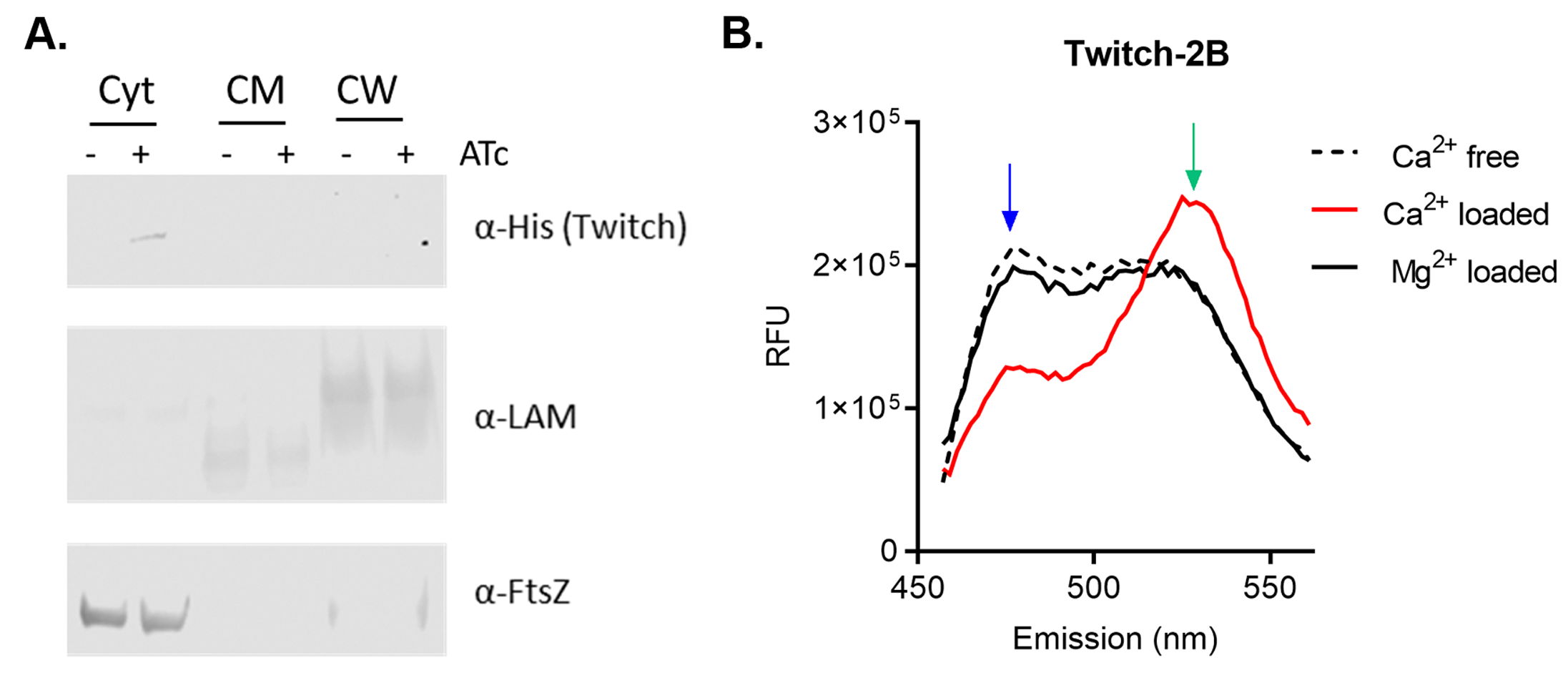

### Supplementary Figure S3

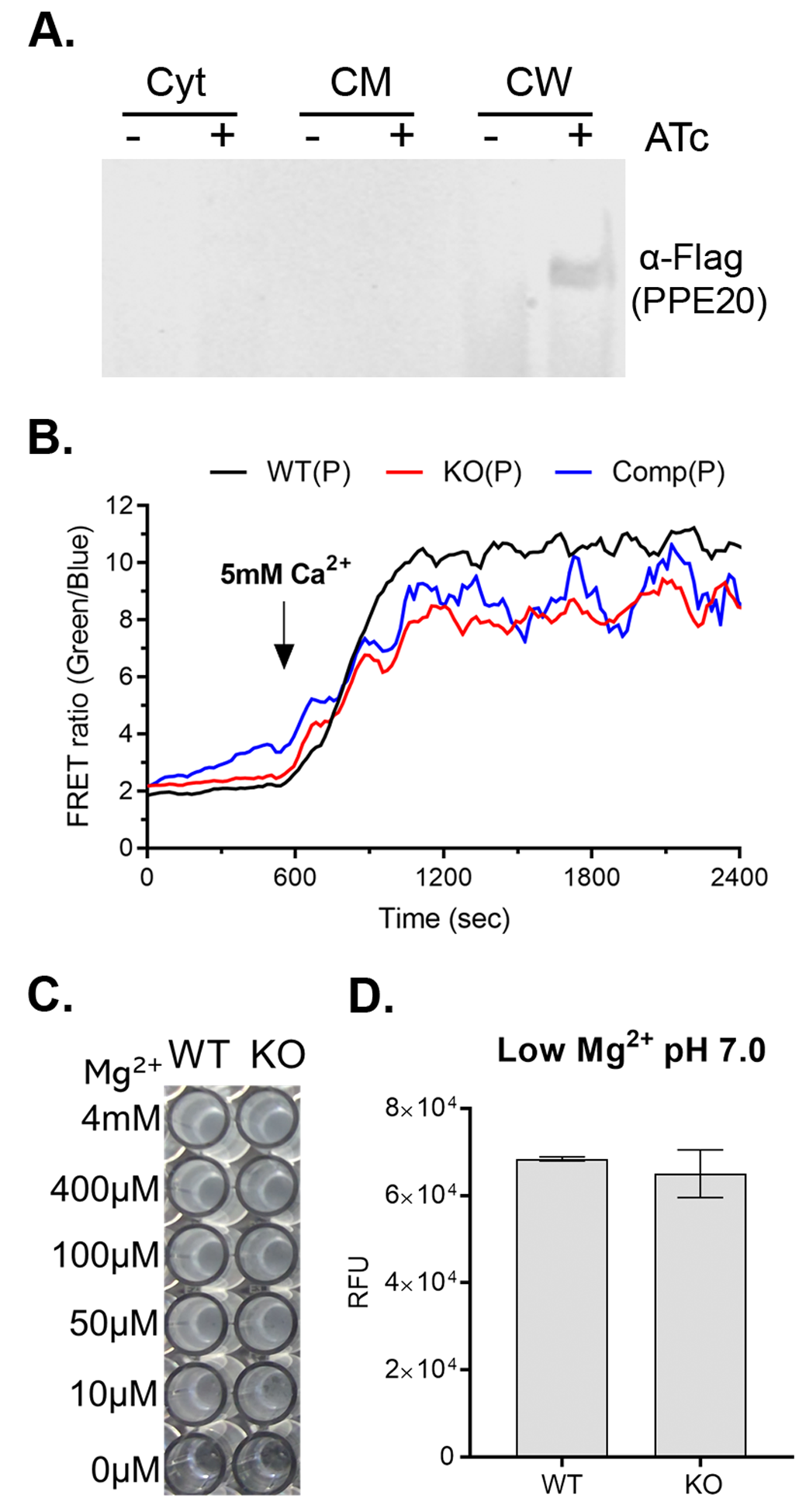
